# Combining Digital Imaging and Genome Wide Association Mapping to Dissect Uncharacterized Traits in Plant/Pathogen Interactions

**DOI:** 10.1101/296939

**Authors:** Rachel F. Fordyce, Nicole E. Soltis, Celine Caseys, Raoni Gwinner, Jason A. Corwin, Susana Atwell, Daniel Copeland, Julie Feusier, Anushriya Subedy, Robert Eshbaugh, Daniel J. Kliebenstein

## Abstract

Plant resistance to generalist pathogens with broad host ranges, such as *Botrytis cinerea*, is typically quantitative and highly polygenic. Recent studies have begun to elucidate the molecular genetic basis underpinning plant-pathogen interactions using commonly measured traits including lesion size and/or pathogen biomass. Yet with the advent of digital imaging and phenomics, there are a large number of additional resistance traits available to study quantitative resistance. In this study, we used high-throughput digital imaging analysis to investigate previously uncharacterized visual traits of plant-pathogen interactions related disease resistance using the *Arabidopsis thaliana/Botrytis cinerea* pathosystem. Using a large collection of 75 visual traits collected from every lesion, we focused on lesion color, lesion shape, and lesion size, to test how these aspects of the interaction are genetically related. Using genome wide association (GWA) mapping in *A. thaliana*, we show that lesion color and shape are genetically separable traits associated with plant-disease resistance. Using defined mutants in 23 candidate genes from the GWA mapping, we could identify and show that novel loci associated with each different plant-pathogen interaction trait, which expands our understanding of the functional mechanisms driving plant disease resistance.

**Summary:** Digital imaging allows the identification of genes controlling novel lesion traits.

## Introduction

The ability to resist biotic attackers, including plant pathogens, is central to a plant’s survival and fitness. This need for resistance is complicated by the immense variety of infection strategies and pathogenic species that can attack an individual plant. These pathogens differ in their lifestyle and host range with each having a different requisite optimal virulence mechanism. Specialist pathogens often co-evolve with host plants to develop virulence strategies targeted on a specific resistance mechanisms within the plant. These biotrophic pathogens must strike a delicate balance between keeping host plant’s cells alive by while absorbing nutrients and avoiding detection and suppressing plant defenses. Plant defense against specialists is accordingly tailored to the specific pathogen with which it has been co-evolving, often involving gene products that directly or indirectly recognize effectors or molecular patterns specific to that pathogen. This co-evolutionary link between host and specialist pathogen frequently leads to large-effect, gene-for-gene resistance involving the specific pathogen recognition R-genes (Jones and Dangl, 2006). In contrast, necrotrophic pathogens are often generalists, deploying a diverse array of virulence mechanisms to precisely kill host plant cells (Laluk and Mengiste, 2010). The generalist attack is multifaceted, relying on many different phytotoxins and enzymes working in parallel to subjugate the plant. Host-pathogen co-evolution in generalist pathogens is not well understood because the pathogen can attack innumerable hosts, causing myriad plant species to encounter a similarly dizzying array of generalist pathogen virulence mechanisms. As such, generalist necrotrophic resistance typically has a highly quantitative polygenic basis that involves physical barriers, such cuticle formation and cell wall modification, chemical barriers, such as secondary metabolites or reactive oxygen species, and inducible defenses (Rowe and Kliebenstein, 2008; Laluk and Mengiste, 2010; Windram et al., 2012; Corwin et al., 2016b).

Infected plants display a large variety of interrelated traits during the infection process. The abundance of scientific literature concerning molecular mechanistic analysis of plant-pathogen interactions largely relies either on lesion size (a single number measuring the plant area displaying symptoms) or pathogen biomass within the infected plant tissue. While these observations are relatively easily measured traits that reflect plant susceptibility to a pathogen, they do not fully describe all aspects of disease resistance within the plant host. For instance, comparing the molecular and quantitative genetic basis of resistance using lesion size versus the underlying biochemical plant resistance responses has shown that different genes can contribute to these traits. This suggests that the lesion size measurements are not providing the full picture of the resistance response (Rowe and Kliebenstein, 2008; Bock et al., 2010; Li et al., 2015; Corwin et al., 2016b; Schwanck and Del Ponte, 2016; Barbedo, 2017; Matsunaga et al., 2017). There has been recent interest in extending our understanding of plant-pathogen resistance by conducting more extensive phenotyping of disease symptoms, including hyperspectral imaging of lesions that records spectra from the visible into the infrared range (Montes et al., 2007; Kuska et al., 2015; Mutka and Bart, 2015; Leucker et al., 2016). This analysis has differentiated between diseases on the same plant through biochemical responses using light spectra from 400nm to 1000nm (Mahlein et al., 2012). These spectra provide a blend of visual symptoms characterized not only by extent of necrosis, but also by water content, pigment modification, tissue function, and appearance of fungal structures (Mahlein et al., 2012). However, the molecular mechanisms controlling these visual and biochemical aspects of the plant-pathogen interaction are largely unstudied, and little is known about their mechanistic basis and how this relates to molecular mechanisms identified using pathogen biomass or lesion size. Therefore, there is a need to study plant-pathogen interactions from the multi-dimensional view that can be provided by digital imaging of lesions and assess how this links to one-dimensional measurement of lesion progression. This could discover new insights into the physiology of pathogen strategy and host response.

One developing model system to directly study the complexity of molecular plant resistance to generalist pathogens can be found within the *Arabidopsis thaliana/Botrytis cinerea* pathosystem (Corwin and Kliebenstein, 2017). *Botrytis cinerea* is a generalist necrotrophic pathogen with a broad range of hosts that includes Bryophytes, through Gymnosperms, to Angiosperms (Jarvis, 1977; Staats et al., 2005; Choquer et al., 2007; Elad et al., 2016). This host range appears to be facilitated by a high level of intraspecific genetic diversity and elevated levels of recombination in comparison to specialist clonal pathogens (Giraud et al., 1997; Alfonso et al., 2000; Vaczy et al., 2008; Kretschmer et al., 2009; Stefanato et al., 2009; Amselem et al., 2011; Aguileta et al., 2012; Atwell et al., 2015). This natural genetic variation within *B. cinerea* quantitatively affects lesion size within all plants tested, including *A. thaliana* (Rowe and Kliebenstein, 2010; Valiuskaite et al., 2010; Amselem et al., 2011; Corwin et al., 2016a). *A. thaliana* resistance to *B. cinerea* involves key signaling pathways, such as Jasmonate and the BIK/BAK response system (Thomma et al., 1999; Veronese et al., 2006). More recently, high-throughput digital imaging analysis of the *A. thaliana/B. cinerea* interaction has allowed the identification hundreds of genes associated with the size of the lesion, with most genes being associated to one or two isolates (Corwin et al., 2016a). The high-throughput digital imaging pipeline used also can provide additional visual traits that facilitates the comparison of novel visual traits with traditional measures related to disease resistance. We examined a wide array of visual information of lesion traits to find a broad and largely orthogonal subset of visual parameters to measure the interaction, including lesion yellowness, lesion greenness, and lesion shape.

Here, we rely on the natural variation in the host and pathogen for the Arabidopsis/Botrytis pathosystem to measure a variety of novel infection-related traits and to conduct genome wide association (GWA) mapping within the host. Genetic mapping will allow us to identify host genes associated with these novel visual traits and expand our understanding of the genetic architecture of quantitative resistance against generalist pathogens. Previous work has shown that GWA mapping, when applied to the *Arabidopsis thaliana/Botrytis cinerea* pathosystem, can identify genes involved in polygenic susceptibility traits, such as lesion size and camalexin production (Corwin et al. 2016). This approach also does not impart *a priori* assumptions on the genes or mechanisms that may play a role in controlling the trait in question. This is exemplified by the observation that many of the genes identified by GWA mapping in *A. thaliana* have not been previously linked to this pathosystem (Jarvis, 1977; Staats et al., 2005; Choquer et al., 2007; Rowe and Kliebenstein, 2008; Mengiste et al., 2010; Valiuskaite et al., 2010; Amselem et al., 2011; Corwin et al., 2016b; Corwin and Kliebenstein, 2017). For example, this identified Late-elongating Hypocotyl (*LHY*; *At1g01060*), a key developmental and circadian-regulated gene not previously linked to disease resistance as having a significant effect *on B. cinerea* resistance (Corwin et al., 2016b). In addition to known genes, this approach could identify disease resistance effects in completely novel genes like At4g01860, an uncharacterized Cul4-RING E3 component. This suggests that extending this analysis to other traits may allow the identification of additional new disease resistance loci. Further, this can be directly facilitated by a reanalysis of the digital images collected from the previous study explore other traits related to disease resistance.

Within the current study, we re-analyzed the images collected from our previous work to measure visual aspects of lesion development that are not commonly measured with respect to disease resistance, specifically lesion shape and color. A genome wide association mapping analysis of select traits in Arabidopsis revealed a large compendium of genes associated with lesion size, lesion yellowness, lesion greenness, and lesion shape. GO enrichment analysis showed that there were biological processes specific to subsets of traits rather than processes affecting all of the lesion-associated traits. To validate the results of the GWA mapping, we selected 23 genes associated with one or more of the lesion traits and compared the phenotypes of T-DNA knockout mutants. The validation rate was highest for lesion size, at 60%, and lower for the color and shape traits: 33% for lesion yellowness, 38% for lesion greenness, and 20% for lesion eccentricity. This further demonstrated that the genes affected subsets of lesion traits rather than being specific to all traits. As such, expanding our phenotypic analysis of plant-pathogen interactions to novel visual traits like lesion shape and color can identify different mechanisms than solely focusing on lesion size.

## Results

### Identification of Alternative Disease Related Traits

Using a custom digital image analysis pipeline, we had previously measured lesion size (area) to identify *A. thaliana* genes controlling lesion size within the *Arabidopsis thaliana/Botrytis cinerea* pathosystem using four diverse pathogen genotypes (Corwin et al., 2016b). This previous study used 96 diverse *A. thaliana* accessions chosen based on similar flowering time to minimize ontogenic variation, which also decreased population structure and rare alleles within the collection. The host pathogen interaction was digitally recorded within a randomized complete block design over two experiments with 4 independent biological replicates per experiment per genotype. Specific lesion traits were obtained using a digital image analysis pipeline for which we have previously only reported results for lesion size (Corwin et al., 2016b). Using the same images, we adapted our imaging pipeline to quantify 75 different lesion parameters from the infections. These included a wide range of traits from the number of pixels for specific hues within the lesion, lesion perimeter, proportion of leaf the lesion occupied, length of the major and minor axes of the lesion, eccentricity of the perimeter, etc. (Corwin et al., 2016a)(Figure 1). As previously reported, there is a wide range of lesion sizes across the host/pathogen interactions (Figure 1A versus C). In addition to lesion size, there is variation in color both within the lesion and the residual leaf. For example, some host/pathogen interactions have large chlorotic sectors surrounding the lesions (Figure 1A, C and F), while others have less chlorosis surrounding the developing lesion (Figure 1B and E). In previous work, we have shown that the pathogen is limited solely to the developing lesion and these chlorotic regions are systemic plant responses to the infection (Rowe and Kliebenstein, 2008; Corwin et al., 2016a). Even within the lesion there are color variants with some being yellower (Figure 1A) while others have a greener hue (Figure 1D). In addition to lesion color, there is variation in lesion shape with some lesions being highly symmetrical and circular (Figure 1D and F) while others are elongated along the major axis of the leaf, possibly due to extended pathogen growth along the mid-vein (Figure 1A and B). This preferential mid-vein growth has also been previously seen as genetically variable across *B. cinerea* infecting *A. thaliana* (Corwin et al., 2016a). We proceeded to study if these additional measurements of the interaction between the plant and the pathogen were providing unique genetic insight into the virulence outcome that was not readily obtained from solely lesion size.

**FIGURE 1.**
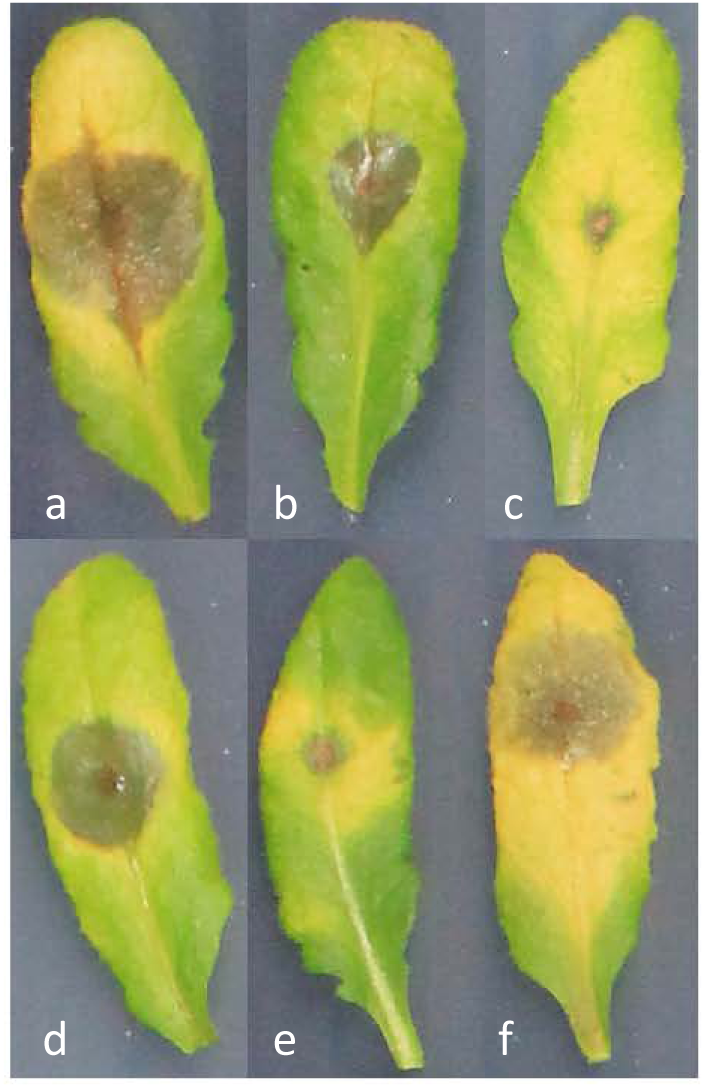
Illustration of different lesion traits. The six *A. thaliana* leaves infected with *B cinerea* represent differing combinations of lesion traits observed. Lesions can be more circular or more eccentric, with pointed ends growing along the mid-vein of the leaf. Greenness and yellowness of both within the lesion and surrounding tissue is also highly variable. The water soaked greyish sectors represent the actual infected lesion. a. Large eccentric lesion with enhanced senescence b. Moderate eccentric lesion with low senescence c. Small lesion with large senescence d. Moderate circular lesion with moderate senescence e. Small lesion with moderate senescence f. Moderate circular lesion with large senescence

### Hierarchical Clustering Analysis for Trait Selection

To select a subset of traits to study further, we conducted a hierarchical clustering analysis (HCA) and constructed a heat map of the 75 lesion traits across the 96 *A. thaliana* accessions infected with the different *B. cinerea* isolates (Figure 2). This analysis allowed us to assess the inter-relatedness of these alternate infection traits both with each other, and with lesion size, to determine whether the traits are providing distinct information with respect to each other. The HCA showed extensive variation in all the traits dependent on *A. thaliana* accessions and *B. cinerea* isolates, leading to four major trait clusters (Figure S1). In these clusters, color and shape traits distinctly clustered apart from lesion size, and varied across plant accession. From this plot, we chose to proceed with three distinct traits for further analysis; lesion yellowness (the proportion of yellow pixels within the lesion to total pixels within the lesion), lesion greenness (total mm^2^ of green pixels within the lesion), and lesion eccentricity (maximum lesion diameter – minimum diameter/maximum lesion diameter). These traits, along with lesion size, represent a broad sampling of the identified variation in the traits and provide a diverse array of measurable aspects of the *A. thaliana – B. cinerea* interaction.

**FIGURE 2.**
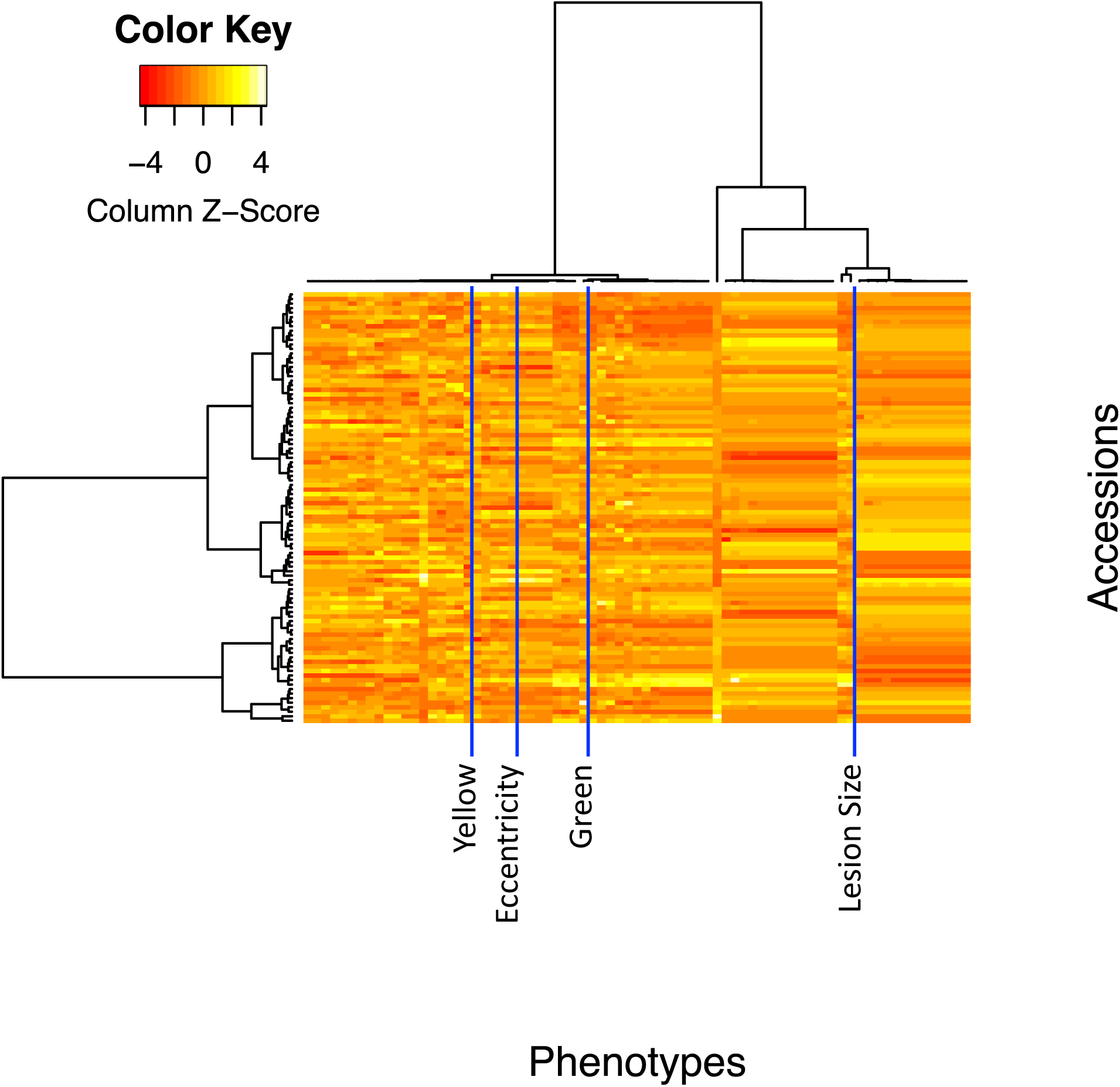
Hierarchical cluster analysis of lesion traits. The heat map shows the collection of 75 lesion measurements of the four *B. cinerea* isolates (columns) on the collection of 96 *A. thaliana* accessions (rows). Labeled lines show lesion size, yellowness, greenness and eccentricity for the *B. cinerea* BO5.10 isolate.

From the HCA, lesion yellowness provides a trait that is distinct from lesion size suggesting that it may provide a highly unique measure of the interaction. We also focused on lesion greenness to increase our sampling within the hierarchical cluster space around lesion yellowness. Lesion greenness may be related to the presence of green islands that were noted in other plant pathogen interactions but have not been commonly studied (Walters et al., 2008). Lesion eccentricity was chosen as it is a spatial trait that is uncorrelated with lesion size traits. Previous work in our lab had suggested that lesion shape variation may be linked to genetic variation in the pathogens ability to identify and grow directionally along the mid-vein of the host leaf (Corwin et al., 2016a). Thus, including this trait may provide unique genetic insight into the ability of the pathogen to directionally orient growth along primary leaf vasculature within the host. We also included lesion size (mm^2^) in our analysis to provide a direct comparison to previous reports. Thus, these four traits appear to have natural genetic variation both within the plant and pathogen that may provide unique insights into the plant/pathogen interaction.

### Genetic Variation in New Lesion Traits

We utilized the phenotypic measurements for all four traits across all genotypic interactions of *A. thaliana* and *B. cinerea* to quantitatively assess the role of host and pathogen genetic variation in controlling these traits. We used linear models to analyze the variance across the 96 *A. thaliana* accessions and the four *B. cinerea* isolates. The genetic variation in both the pathogen and the host significantly influenced all four resistance associated traits (Table1). The proportion of variance attributable to *A. thaliana* accession was significant for these four traits, although relatively small for this experiment. The heritability across the entire dataset that was solely attributable to *A. thaliana* was relatively low albeit significant for each trait tested (Table 1). The proportion of heritability attributable to pathogen genetic variation was higher for the color and size traits ranging from 10-40% but there was no significant heritability within this small collection of isolates ascribed to pathogen variation for lesion eccentricity (Table 1). In contrast, the eccentricity was controlled by the interaction of genetic variation in the host and pathogen (Table 1), suggesting that this trait is highly dependent on the interaction between the pathogen genotypes and the different plant accessions. Thus, there is significant genetic variation for all traits that is dependent upon both the host and pathogen genomic diversity.

**TABLE 1.** Heritability of Lesion Traits. For each of the four traits, the table displays the sum of squares for each term in an analysis of variance, p values for each term, and the calculated heritability (proportion of total variance) attributed to the specific model terms. The analysis used the model Lesion Trait ~ Experiment + Accession * Isolate.

|  | SS | P | Heritability |
| --- | --- | --- | --- |
| **Size** |  |  |  |
| Experiment | 1224 | 0.002 |  |
| Plant | 179962 | < 0.001 |  |
| Accession | 25273 | < 0.001 | 3.0% |
| Isolate | 333466 | < 0.001 | 39.5% |
| Accession:Isolate | 57552 | < 0.001 | 6.8% |
| **Yellow** |  |  |  |
| Experiment | 2387113 | 0.036 |  |
| Plant | 964232385 | < 0.001 |  |
| Accession | 70751623 | 0.014 | 2.5% |
| Isolate | 557388090 | < 0.001 | 20.0% |
| Accession:Isolate | 176639269 | 0.083 | 6.3% |
| **Green** |  |  |  |
| Experiment | 3 | 0.281 |  |
| Plant | 3958 | < 0.001 |  |
| Accession | 756 | < 0.001 | 6.1% |
| Isolate | 1232 | < 0.001 | 10.0% |
| Accession:Isolate | 1084 | < 0.001 | 8.8% |
| **Eccentricity** |  |  |  |
| Experiment | 0.00070 | 0.842 |  |
| Plant | 13.04996 | 0.003 |  |
| Accession | 2.34077 | 0.008 | 4.3% |
| Isolate | 0.00387 | 0.974 | <0.1% |
| Accession:Isolate | 6.00843 | 0.022 | 11.1% |

For further analysis, we obtained the model-corrected least squared means from these linear models to account for the environmental influence. This should help to focus the analysis on the genetic influence of the plant host in the downstream genetic mapping. Given the significance of the interaction between plant accession and pathogen isolate, we analyzed each lesion trait individually for each pathogen genotype across the diverse *A. thaliana* accessions as a separate trait to maximize our ability to identify causal loci.

### Analysis of Covariance of Lesion Traits

To test how the lesion size, color, and shape traits may correlate across the different host/pathogen combinations, we conducted an ANCOVA (Analysis of CoVariance) comparing each pair of traits across the 96 diverse *A. thaliana* accessions for each of the four *B. cinerea* isolates (Figure 3). Significant correlations (p <0.05) between the color and size, traits indicate that these three traits are generally related. Lesion yellowness and greenness show a positive correlation, both overall and within each *B. cinerea* isolate group, and this correlation is statistically consistent between groups (Figure 3). In contrast, lesion color (both greenness and yellowness) and size, display significant variation amongst the *B. cinerea* isolates for the relationship with B05.10 and Supersteak isolates showing a negative correlation between lesion yellowness and size while Apple517 and UKRazz show a neutral or slightly positive correlation (Figure 3). In contrast to color and lesion size, eccentricity does not display uniform correlations across all the isolates. Instead, the significant correlations are highly dependent on the interaction with isolate (Figure 3). For example in eccentricity by lesion size, there is a strong negative correlation in the *B. cinerea* isolates B05.10 and Supersteak, while only a very slight positive correlation in Apple517 and UKRazz.

**FIGURE 3.**
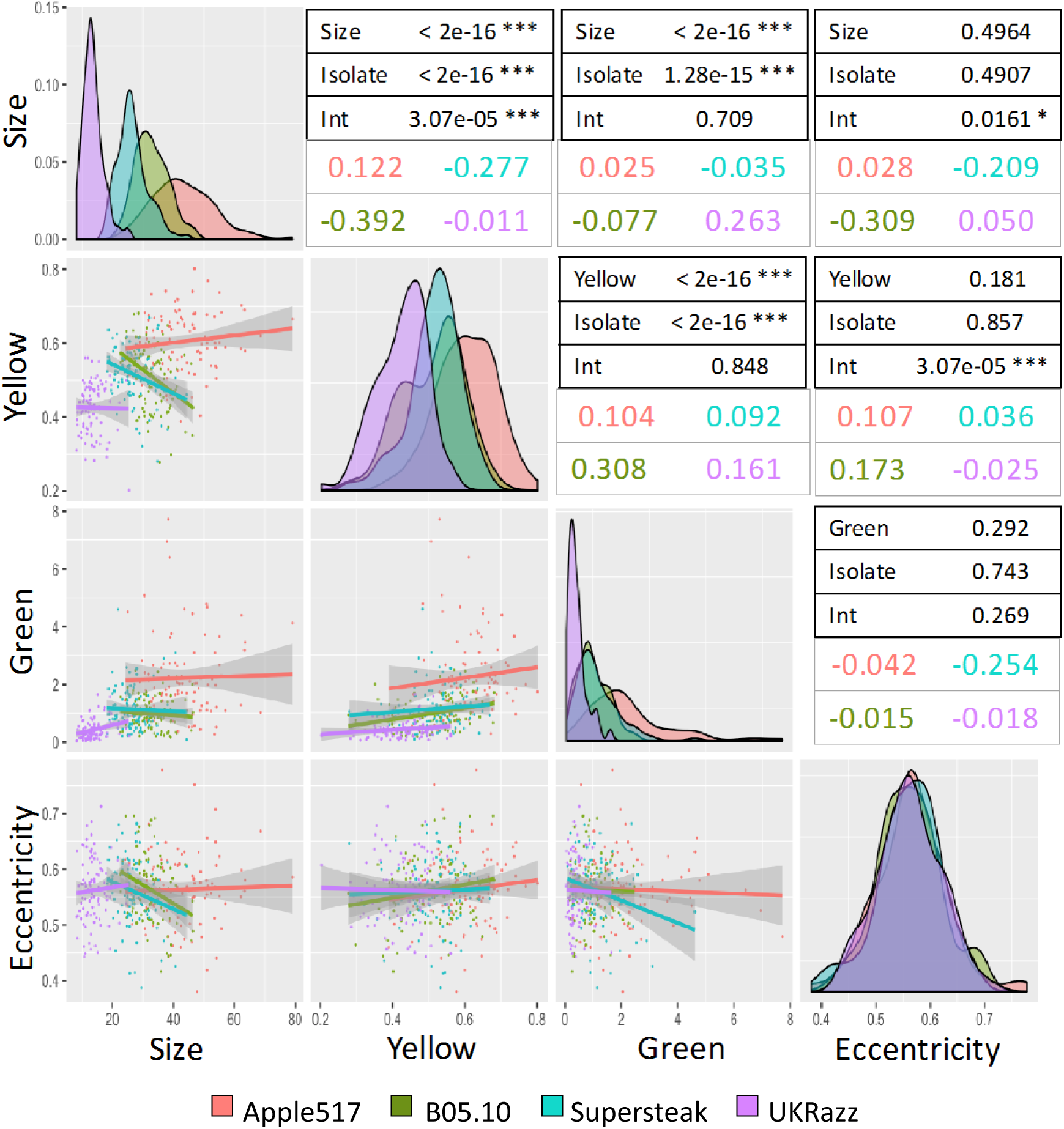
Isolate dependency of Ttait correlations across the isolates. ANCOVA was used to test for correlations between each pair of traits across the 96 *A. thaliana* isolates using each of the four different *B. cinerea* isolates. Scatter plots with 95% confidence interval for the trait correlation in each isolate are shown on the left half of the plot. The diagonal shows the distribution of the specific trait across the four *B. cinerea* isolates. The right hand tables show the p-values for the trait-trait correlations, *B. cinerea* isolate dependency of these correlations, and the interaction of *B. cinerea* isolate on the trait-trait correlation (Int) using the following model). The estimated slopes for each trait-trait interaction for each *B. cinerea* isolate are shown by the colored text.

This suggests that while there is some common and some trait dependent genetic variation in how *A. thaliana* responds to these diverse *B. cinerea* isolates using these lesion traits. This analysis revealed that while there is some correlation between the four lesion traits, the host and pathogen genotypes alter these relationships indicating that there is not a simple mechanistic basis for these relationships. Further, the four lesion traits represent measures of distinct and nuanced interactions between the plant and pathogen, and that the genes associated with each trait likely influence different mechanisms in a multifaceted genetic storyline.

### Genome-Wide Association Mapping of Lesion Traits

To identify *A. thaliana* genes that are associated with these new lesion traits, we conducted a genome-wide association (GWA) study. We used the model-corrected trait means for each *A. thaliana* accession infected independently with the four different *B. cinerea* isolates across the traits of lesion size, eccentricity (shape), and yellow and green color. We employed a publicly available database of genomic polymorphisms across the 96 *A. thaliana* accessions consisting of 115,301 SNPs with minor allele frequency (MAF) of > 0.2 (Atwell et al., 2010; Corwin et al., 2016b). The model for GWA mapping used a previously published ridge regression approach with a permutated-effects threshold, which has previously been used to successfully identify causal loci for both plant/pathogen interactions and plant metabolic variation (Shen et al., 2013; Corwin et al., 2016b; Francisco et al., 2016; Kooke et al., 2016). Using the means for each *A. thaliana* accession for each of the four lesion traits in response to the four individual *B. cinerea* isolates, we conducted GWA mapping for each combination separately for each isolate, resulting in four separate GWA maps for each trait. Because the variance and mean of each lesion phenotype differed according to pathogen genotype, the resulting effect sizes for *A. thaliana* SNPs also varied in magnitude. To create comparative Manhattan plots of the results for a single trait across all four *B. cinerea* isolates, we z-scaled the SNP effect sizes within each pathogen and overlaid the results (Figures 4 and 5). We calculated an effect size significance threshold for each individual trait by permuting the data one thousand times and running the ridge regression on the permuted data, and locating the 95^th^ percentile (Tables S3-S6).

**FIGURE 4.**
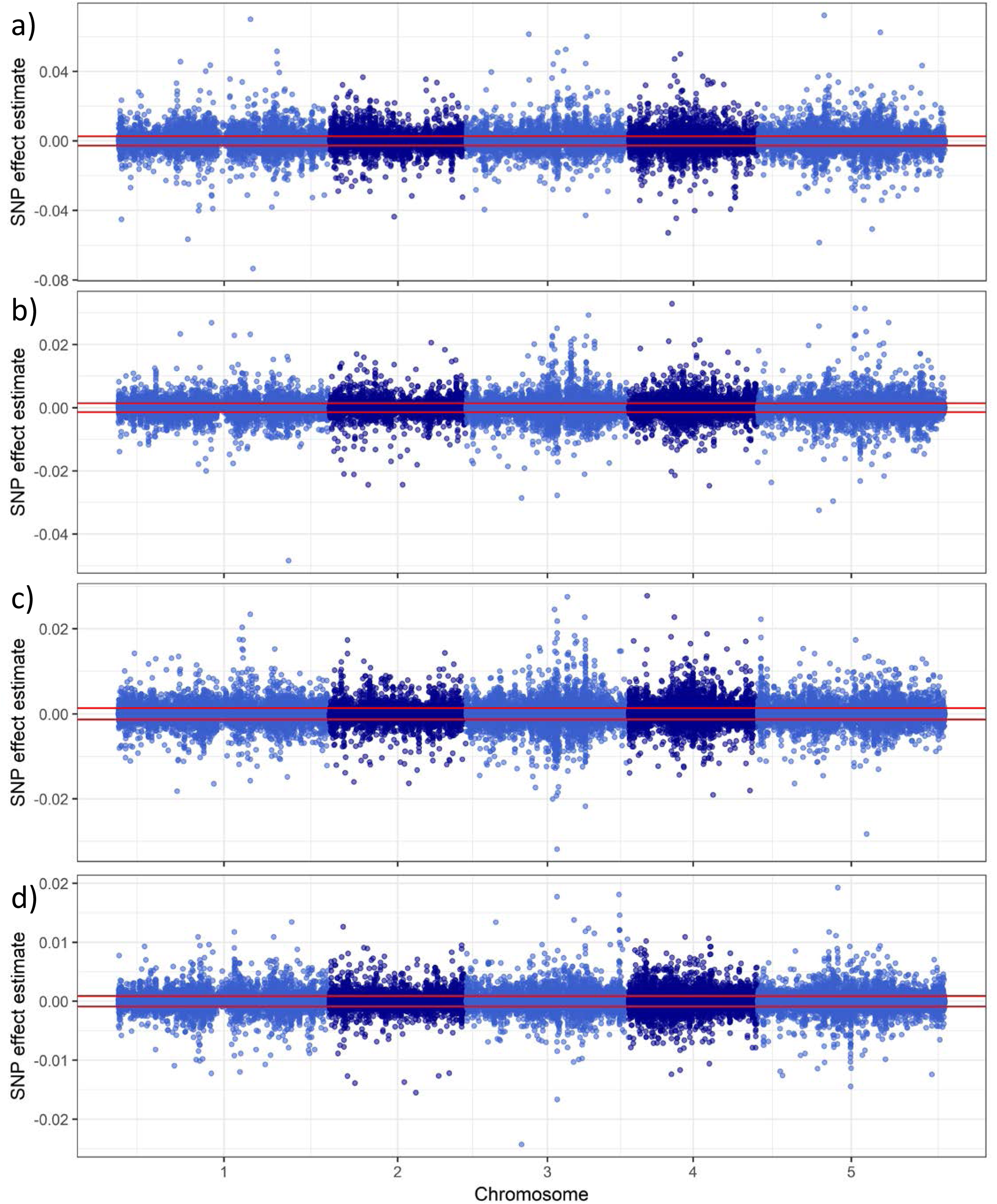
Lesion size GWAS for four *B. cinerea* isolates. Manhattan plots showing lesion size trait GWAS results, as measured on four *B. cinerea* isolates: a, Apple517; b, B05.10; c, Supersteak; and d, UKRazz. The horizontal red-line shows the significance threshold as estimated by permutation.

**FIGURE 5.**
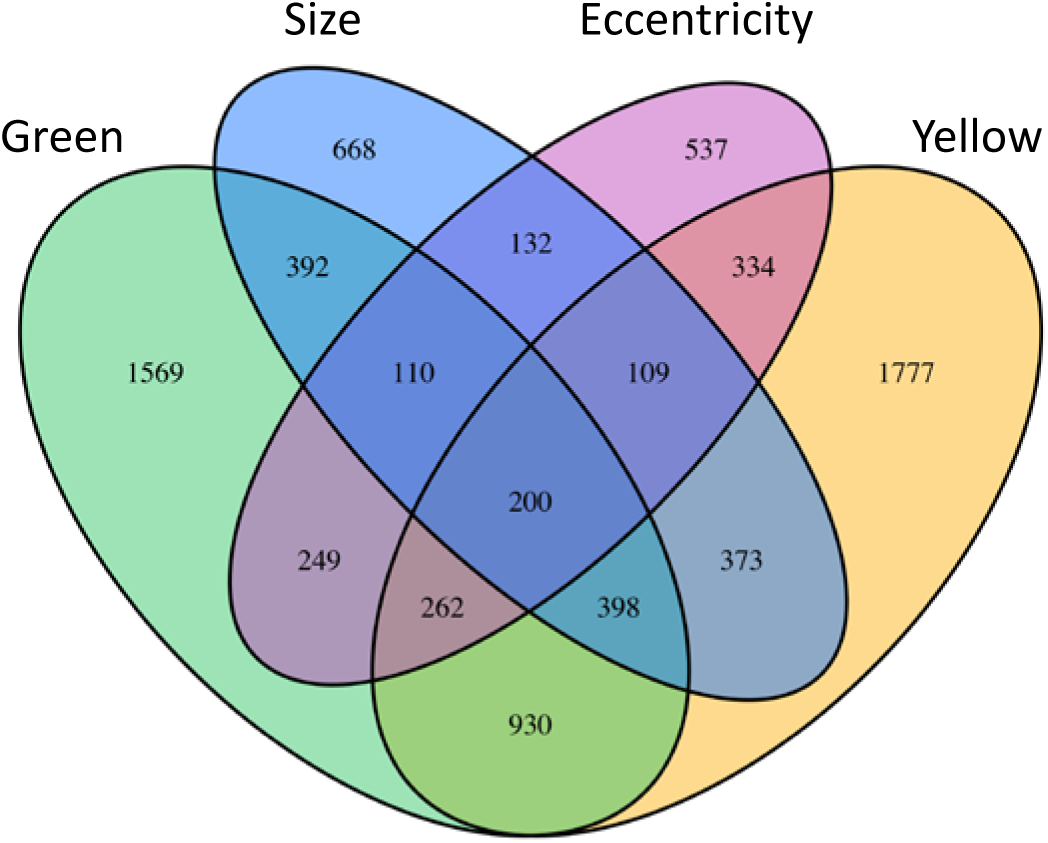
Venn diagram of genes found in GWA mapping. A Venn diagram of the candidate *A. thaliana* genes found to associate with each lesion trait via GWA. The number of genes listed for each lesion trait reflects the number of genes with two or more SNPs above the 95^th^ percentile threshold for that particular trait.

The comparative Manhattan plots (Figures 4, S2 and S3) showed that all four lesion associated traits lacked large, distinct SNP peaks. Instead, there are numerous significant SNPs with small-to-medium effect sizes distributed across the genome, suggesting a highly polygenic nature underlying the variation in these traits. Further, the significant SNPs different between the *B. cinerea* isolates, suggesting that most SNPs are dependent upon the interaction between plant and pathogen genotypes (Figures 4 and 5). For lesion Yellowness and Eccentricity, only Apple517 had significant SNPs possibly because this isolate had the highest level of phenotypic diversity in these traits (Figure 3). We next proceeded to identify putative causal loci in the GWA results by calling a gene as significantly associated with a trait if two or more SNPs within the coding region and 1 kb of the 5’ upstream and 3’ downstream regions had effect sizes above the 95^th^ percentile threshold (Chan et al., 2011; Corwin et al., 2016b). This approach has previously been shown to reduce false positives within the dataset (Corwin et al., 2016b). Combining the analyses of each trait across the *B. cinerea* isolates for which there were significant SNPs identified 2,382 genes associate with lesion size, 4,383 with lesion yellowness, 4,110 with lesion greenness, and 1,933 with lesion eccentricity. Agreeing with the hypothesis that these traits can identify new cellular mechanisms, the majority of the genes identified are associated with only one or a few traits (Figure 5). Few genes (2.5% of 7940 total genes identified) were associated with all four lesion related traits (Figure 5). Of the genes associated with lesion size, 8% were not associated with any of the other three lesion traits. Likewise, 22% of genes associated with yellowness were associated exclusively with that phenotype, 20% for green, and 7% for eccentricity. Overall, 57% of the genes found in this study associated with only one of the lesion traits and 31% associate with exactly two of the lesion traits. These results reinforce the idea that while there is some correlation between the traits, they are quantitative traits that represent different measures of the plant-pathogen interaction.

### GO Category Enrichment Analysis

We performed a gene ontology (GO) category enrichment analysis on biological processes of genes associated with each of the four traits to identify *A. thaliana* mechanisms that may influence variation in lesion traits. We performed GO enrichment analysis for each set of candidate genes (Table 2). Both lesion color traits linked to genes involved in defense against pathogens and insects (Table 2). They also identified unique enrichment categories wherein each trait had different enriched categories. Specifically, lesion greenness shows enrichment in maintenance of shoot apical meristem identity, and lesion yellowness in lignin biosynthetic process. In contrast, eccentricity associated candidate genes are enriched in a variety of unexpected processes, such as negative regulation of flower development and negative regulation of reproductive process, in addition to known pathogen-associated processes, such as cell wall thickening and callose deposition in cell walls. Genes controlling cell wall modification play a role in plant defense strategies to contain the infected area, but have not been studied in the context of lesion shape (Cantu et al., 2008; Mengiste et al., 2010; Bethke et al., 2016). Thus, these novel visual traits are identifying known defense mechanisms while also extending analysis out to new pathways.

**TABLE 2.** GO Categories enrichment. The table displays the top ten enrichment categories (biological processes) associated with each of the four lesion traits.

| GO.ID | Term | p | Trait |
| --- | --- | --- | --- |
| GO:0006952 | defense response | 1.40E-05 | Size |
| GO:1901264 | carbohydrate derivative transport | 0.0003 | Size |
| GO:0009617 | response to bacterium | 0.0007 | Size |
| GO:0006749 | glutathione metabolic process | 0.0027 | Size |
| GO:0009556 | microsporogenesis | 0.0028 | Size |
| GO:0006897 | endocytosis | 0.0043 | Size |
| GO:0030029 | actin filament-based process | 0.0048 | Size |
| GO:0048236 | plant-type spore development | 0.0050 | Size |
| GO:0072523 | purine-containing compound catabolic process | 0.0061 | Size |
| GO:1903409 | reactive oxygen species biosynthetic process | 0.0081 | Size |
| GO:0009910 | negative regulation of flower development | 0.0011 | Eccentricity |
| GO:2000242 | negative regulation of reproductive process | 0.0015 | Eccentricity |
| GO:0006349 | regulation of gene expression by genetic imprinting | 0.0021 | Eccentricity |
| GO:0071514 | genetic imprinting | 0.0021 | Eccentricity |
| GO:0052386 | cell wall thickening | 0.0055 | Eccentricity |
| GO:0052543 | callose deposition in cell wall | 0.0055 | Eccentricity |
| GO:0009617 | response to bacterium | 0.0093 | Eccentricity |
| GO:0010150 | leaf senescence | 0.0093 | Eccentricity |
| GO:0006821 | chloride transport | 0.0107 | Eccentricity |
| GO:0015700 | arsenite transport | 0.0107 | Eccentricity |
| GO:0006952 | defense response | 0.0002 | Green |
| GO:0010492 | maintenance of shoot apical meristem identity | 0.0046 | Green |
| GO:0002218 | activation of innate immune response | 0.0051 | Green |
| GO:0002253 | activation of immune response | 0.0051 | Green |
| GO:0042742 | defense response to bacterium | 0.00520 | Green |
| GO:0009617 | response to bacterium | 0.0068 | Green |
| GO:0006928 | movement of cell or subcellular component | 0.0092 | Green |
| GO:0030048 | actin filament-based movement | 0.0092 | Green |
| GO:0034614 | cellular response to reactive oxygen species | 0.0093 | Green |
| GO:0048236 | plant-type spore development | 0.0095 | Green |
| GO:0006952 | defense response | 0.0002 | Yellow |
| GO:0016568 | chromatin modification | 0.0011 | Yellow |
| GO:0044699 | single-organism process | 0.0012 | Yellow |
| GO:0009625 | response to insect | 0.0021 | Yellow |
| GO:0044710 | single-organism metabolic process | 0.0022 | Yellow |
| GO:0006325 | chromatin organization | 0.0029 | Yellow |
| GO:0016569 | covalent chromatin modification | 0.0033 | Yellow |
| GO:0009809 | lignin biosynthetic process | 0.0039 | Yellow |
| GO:0042886 | amide transport | 0.0039 | Yellow |
| GO:0006857 | oligopeptide transport | 0.0045 | Yellow |

In contrast to the individual trait candidate gene lists, The top enrichment categories for the 200 *A. thaliana* genes that associate with all four lesion traits were defense response, response to stress, immune response, plant-type hypersensitive response, programmed cell death, chromatin remodeling, and response to salicylic acid (Table S2). Thus, the core genes associated with all four lesion related traits linked in part to known defense genes. Overall, the enrichment categories of genes associated with only a single trait uncovered several distinct and unique biological processes that are tangentially associated with defense while genes shared across four traits were more conservatively associated with defense. These differently represented enrichment categories according to trait reflect varying, non-overlapping plant mechanisms involved in the multifaceted disease resistance against generalists. This analysis revealed four portraits of plant defense that showed some overlap but also revealed the extent of biochemical mechanisms involved in lesion traits.

### Validation of Genes Associated with Lesion Phenotypes

To test whether the trait-associated genes we found in the GWA mapping were causal, we measured the infection-related phenotypes on knockout mutants of genes associated with each trait of interest. We chose 23 genes (Table 3) based on GWA effect-size ranking, specificity or overlap of traits effected, and ready availability of viable homozygous lines of T-DNA knockout mutants. The lines were grown, infected, and phenotyped using the same image analysis software as the initial 95 *A. thaliana* accessions. All mutants were tested for resistance against each isolate using 20 independent biological replicates per genotype spread across two experiments. All the traits were measured on each lesion and each trait was independently tested for a difference between WT and mutant using ANOVA (Table 3). The validation rate for lesion size was 60%, 33% for lesion yellowness, 38% for lesion greenness, and 20% for lesion eccentricity. Interestingly, this correlates with the overall fraction of each trait controlled by genetic variation (Table 1). This suggests that part of the lower validation rates for the new traits may be a power-to-detect effect issue and require increased replication. Another complication is that in our GWA mapping, we observed that effect sizes for individual SNPs within a single gene can be both positive and negative in comparison to the reference Col-0 accession (Figure 6 and Table 3). Thus, the natural loci likely have multiple haplotypes with differing functionalities while the validation mutants are unidirectional loss-of-function mutants within the reference Col-0 accession. We illustrate this with AT4G17010, a component of the transcription factor III B complex, where significantly associated SNPs have both positive and negative effects in comparison to the reference *A. thaliana* accession Col-0 (Figure 6 and Table 3). Three SNPs have effect sizes that fall above the 95^th^ percentile threshold in the positive direction for lesion yellowness, and two have a similarly large magnitude in the negative direction, indicating that the different haplotypes of this gene uniquely affect lesion yellowness (Figure 8). Clustering the accessions according to SNPs within this particular gene separates the haplotypes into five major groups, with groups I and II containing the SNPs with significant negative effect size, groups III and IV containing the SNP of greatest positive effect size, and group V containing the remaining diverse haplotypes (Figure 7). Even in the face of this complication, the T-DNA knockout mutant for the gene AT4G17010, did lead to altered lesion yellowness but not lesion size (Table 3). Thus, we can use GWA from high-throughput digital imaging to identify and validate new genes affecting visual lesion traits beyond lesion size.

**TABLE 3.** Candidate Gene Assessment. The table shows the validation tests on mutants in 23 potential candidate genes for each of the four measures of lesion formation: size, yellowness, greenness and eccentricity. The # of SNPs classified as significant above the 95 percentile threshold for that trait for that gene are shown as well as the significance of the difference between the insertion mutant and WT across all *B. cinerea* isolates (P – Plant) or an interaction with *B. cinerea* isolates (P – Plant:Isolate). Significance was tested using the models mentioned in the Method section. Nominal p-values between 0.1 and 0.05 are provided if at least one other trait showed significance below 0.05.

|  |  |  |  | Size |  |  | Yellow |  |  | Green |  |  | Eccentricity |  |  |
| --- | --- | --- | --- | --- | --- | --- | --- | --- | --- | --- | --- | --- | --- | --- | --- |
| AGI | Gene | Mutant | #SNPs in coding region | # unique SNPs above 95 | P - Plant | P - Plant: Isolate | # unique SNPs above 95 | P - Plant | P - Plant: Isolate | # unique SNPs above 95 | P - Plant | P - Plant: Isolate | # unique SNPs above 95 | P - Plant | P - Plant: Isolate |
| AT2G25540 | CESA10 | cesA10-1 | 11 | 2 | 0.005 | NS | 4 | 0.053 | NS | 2 | NS | NS | 3 | NS | NS |
| AT2G39940 | COI1 | coi1-1 | 4 | 0 | < 0.001 | 0.096 | 4 | < 0.001 | NS | 0 | < 0.001 | NS | 0 | NS | NS |
| AT1G01060 | LHY | lhy-20 | 2 | 2 | 0.006 | 0.093 | 2 | NS | NS | 2 | 0.028 | 0.014 | 1 | NS | NS |
| AT3G26830 | PAD3 | pad3-1 | 1 | 1 | < 0.001 | 0.014 | 0 | 0.004 | NS | 0 | 0.008 | 0.069 | 0 | NS | NS |
| AT3G52430 | PAD4 | pad4-1 | 6 | 0 | 0.002 | 0.062 | 0 | NS | NS | 1 | 0.010 | 0.019 | 4 | NS | NS |
| AT4G39350 | CESA2 | CS818491 | 8 | 3 | 0.020 | NS | 2 | NS | NS | 2 | 0.067 | NS | 0 | 0.090 | NS |
| AT1G22070 | TGA3 | tga3-2 | 1 | 0 | NS | NS | 0 | NS | NS | 0 | NS | NS | 1 | NS | NS |
| AT4G17010 | AT4G17010 | CS861804 | 7 | 3 | NS | NS | 4 | 0.048 | NS | 2 | NS | NS | 1 | 0.020 | NS |
| AT1G14690 | MAP65 | CS872894 | 10 | 0 | NS | NS | 0 | NS | 0.076 | 0 | NS | NS | 0 | NS | NS |
| AT4G01883 | - | CS67496 | 6 | 0 | 0.083 | NS | 0 | NS | NS | 2 | 0.056 | NS | 0 | 0.043 | NS |
| AT4G14420 | AT4G14420 | N508498 | 6 | 2 | 0.046 | NS | 1 | NS | NS | 2 | 0.003 | NS | 2 | NS | NS |
| AT4G14400 | ACD6 | N559132 | 20 | 3 | NS | NS | 5 | NS | NS | 15 | 0.047 | NS | 0 | NS | NS |
| AT4G14368 | - | N574598 | 16 | 3 | NS | NS | 0 | NS | NS | 6 | NS | NS | 0 | NS | NS |
| AT1G31260 | ZIP10 | N604586 | 9 | 5 | NS | NS | 3 | NS | NS | 0 | NS | NS | 6 | NS | NS |
| AT4G01860 | - | N606172 | 13 | 1 | NS | NS | 0 | NS | NS | 6 | NS | NS | 0 | NS | NS |
| AT2G32150 | - | N640067 | 6 | 0 | NS | NS | 0 | NS | NS | 1 | NS | NS | 0 | NS | NS |
| AT4G01880 | - | CS857701<br>N608769 | 4 | 0 | 0.012 | NS | 1 | 0.037 | NS | 1 | 0.079 | NS | 0 | NS | NS |
| AT3G46640 | LUX | lux | 5 | 0 | NS | NS | 0 | NS | NS | 4 | NS | NS | 0 | 0.046 | NS |
| AT3G45780 | PHOT1 | phot1 | 6 | 3 | NS | 0.052 | 1 | NS | NS | 5 | NS | NS | 0 | NS | NS |
| AT4G18130 | PHYE | phyE-2 | 3 | 3 | 0.002 | NS | 0 | 0.024 | NS | 0 | NS | NS | 0 | NS | NS |
| AT4G39030 | SID1 | sid1 | 9 | 0 | NS | 0.014 | 0 | NS | NS | 8 | NS | NS | 0 | NS | NS |
| AT5G67370 | CGLD27 | N529971 | 4 | 0 | NS | NS | 3 | NS | NS | 1 | NS | NS | 0 | NS | NS |
| AT5G45950 | - | N582692 | 5 | 1 | NS | NS | 4 | NS | NS | 0 | NS | NS | 0 | NS | NS |
| AT1G20910 | - | N641443 | 5 | 0 | NS | 0.023 | 0 | NS | NS | 5 | NS | NS | 1 | NS | NS |
| AT5G13550 | SULTR4;1 | N662135 | 5 | 0 | NS | NS | 0 | NS | NS | 0 | NS | NS | 5 | 0.094 | NS |

**FIGURE 6.**
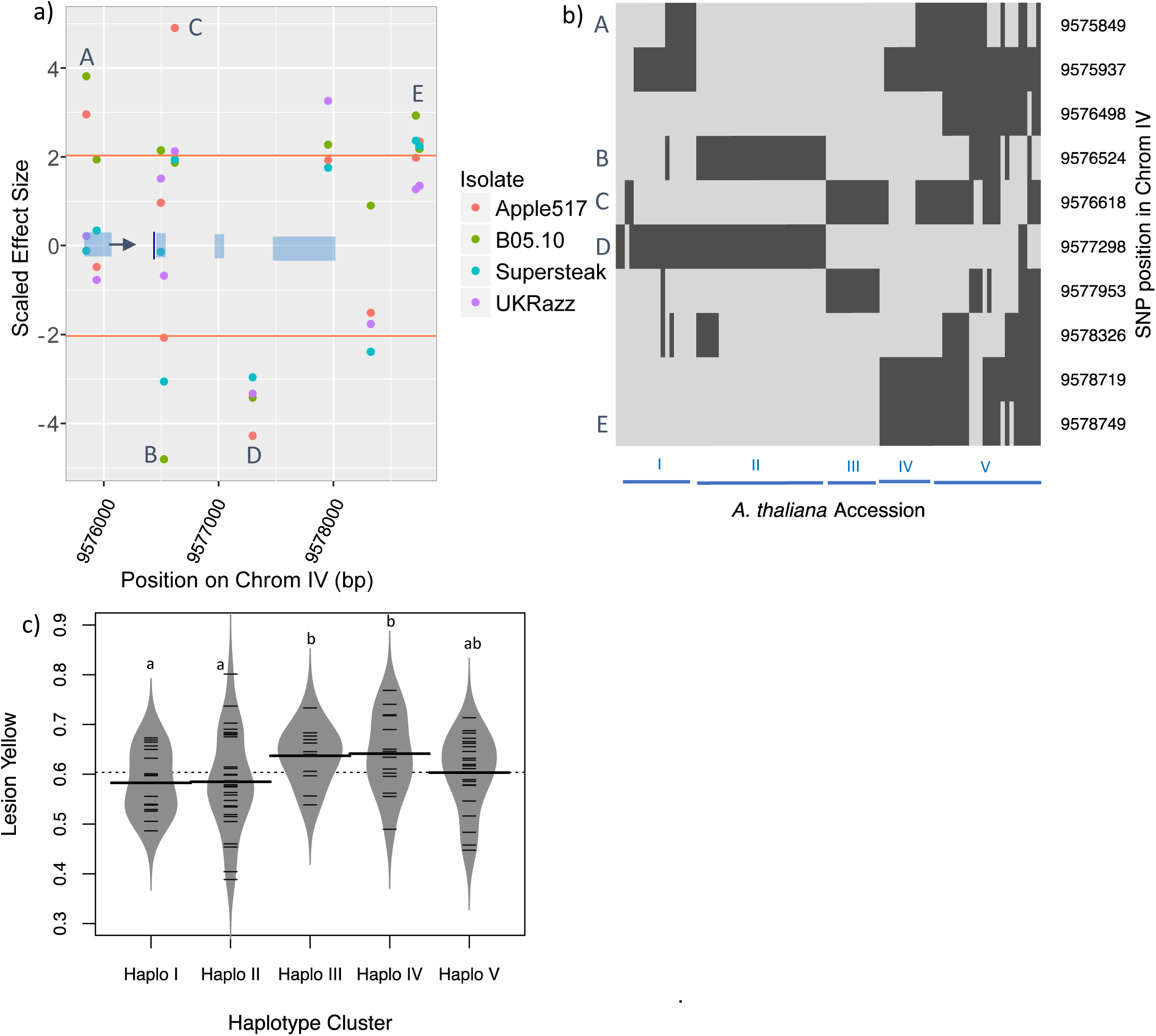
Haplotype diversity effects on trait to genotype linkages using AT4G17010. (a) Plot of z-scaled SNP effect size across all four *B. cinerea* isolates on lesion yellowness within 1000bp of the AT4G17010 coding region (represented in blue bocks) of AT4G17010. The arrow indicates the transcriptional start site. The horizontal orange lines indicate the positive and negative permutation thresholds for the *B. cinerea* isolate Apple517. The letters show the SNPs that are significantly associated with lesion yellowness in *B. cinerea* Apple517. (b) Hierarchical clustering of 95 *A. thaliana* accessions based on SNPs within AT4G17010. Haplotypes are assigned into five major groups, denoted by Roman numerals. Light grey indicates the SNP is the Col-0 allele while dark grey is the opposite allele. The SNPs are presented in their genomic order rather than the haplotype grouped structure. (c) Distributions of lesion yellowness across the *A. thaliana* accessions infected with *B. cinerea* Apple517. The *A. thaliana* accessions are grouped by approximate SNP haplotypes in b. The model corrected mean value of lesion yellowness for *A. thaliana* reference accession Col-0 was 0.545. Significance differences between the groups are shown by letters.

**FIGURE 7.**
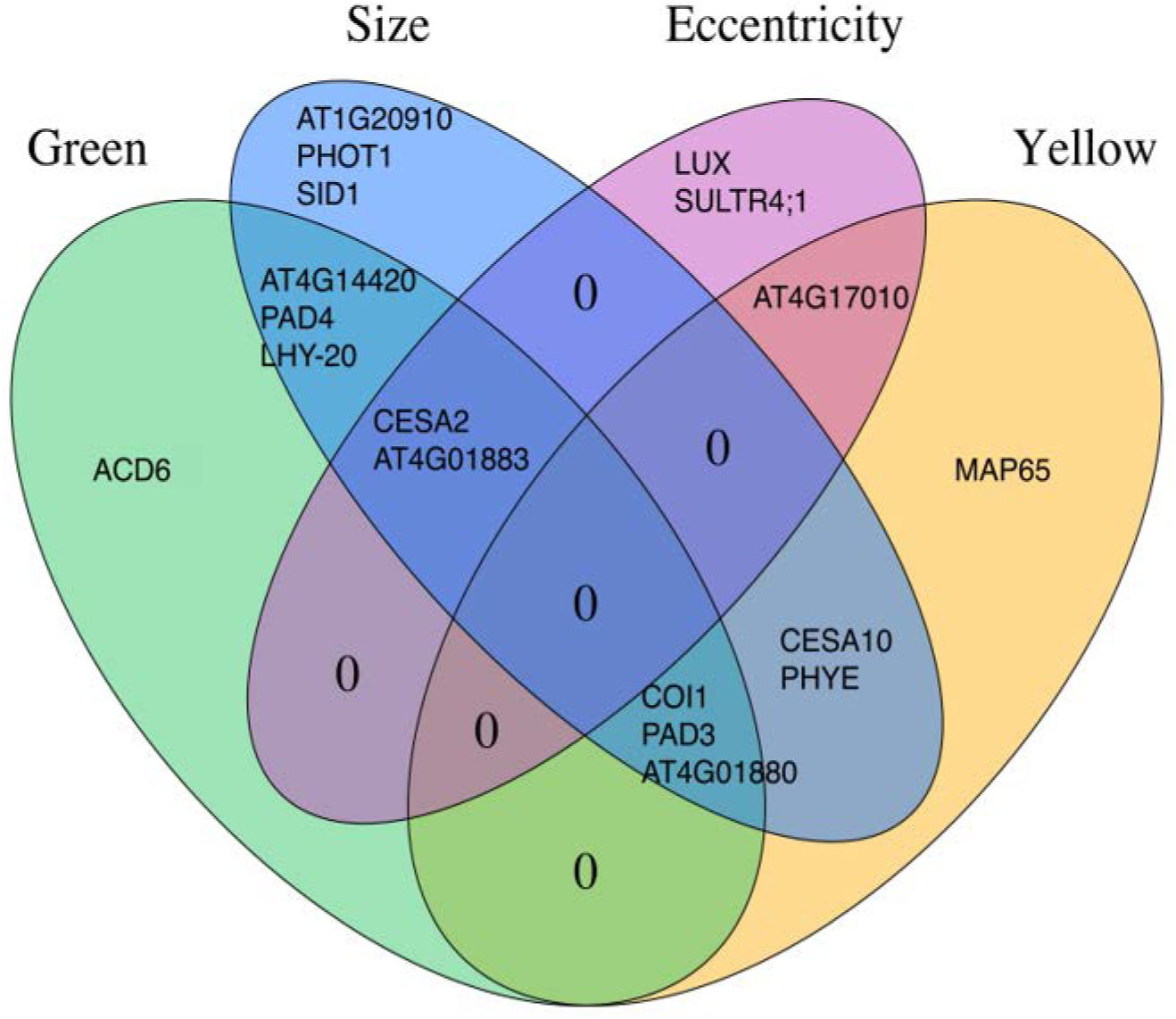
Lesion traits affected by T-DNA insertional mutants. A Venn diagram showing the lesion traits significantly affected by each of the T-DNA knockout mutants in comparison to the wildtype background. The gene identifiers are located within the appropriate section of the diagram showing all the phenotypes altered by mutations in that gene.

### Specific Loci and Trait Overlap

The validation tests showed that the genes were highly specific in their effects on the lesion traits by affecting only one or two traits. Only two genes, CESA2 and AT4G01883, were linked to three traits (Figure 8). Critically, no mutant affected all of the measured traits (Figure 7). This analysis identified new genes not previously linked to resistance to *B. cinerea*. A knockout mutant for *PHYTOCHROME E* (*phyE*), previously linked to shade avoidance, germination, seedling de-etiolation, and flowering time (Halliday and Whitelam, 2003), showed a statistically significant increase in lesion size and lesion yellowness compared to wildtype. It had been previously demonstrated that phyE affects stomatal conductance and photorespiration, which have been shown to play an important role in signaling programmed cell death and systemic acquired resistance defense response (Muhlenbock et al., 2008; Szechynska-Hebda et al., 2010). Other phytochromes, such as phyB, have been linked with defense response in *A. thaliana* via this pathway (Griebel and Zeier, 2008; Zhao et al., 2014) but there is no prior evidence in the literature of phyE affecting resistance. A knockout mutant of *SULFATE TRANSPORTER 4.1 (sultr4;1)* showed a greater lesion eccentricity. Sultr4;1 transports Sulphur from storage in the vacuole, facilitating the synthesis of Sulphur-containing amino acids and glutathione, a compound involved in balancing cellular redox (Zuber et al., 2010b). Sultr4;1 has not been previously linked with pathogen resistance, yet somehow plays a role in the defining the shape of a Botrytis lesion. This gene was neither associated with nor found to affect any of the other lesion traits, highlighting the importance of expanding the visual information used in investigation of lesions. This highlights the utility of measuring different lesion related traits beyond biomass or lesion size to identify new resistance mechanisms.

## Discussion

In the analysis of plant/pathogen interactions, there is a dominant focus on measuring pathogen success either as lesion area or pathogen biomass to quantify the level of resistance/virulence in the interaction. However, this is merely one aspect of the interaction and active infections can affect many different traits within the plant host (Agrios, 2005). New, non-destructive imaging methods are beginning to illuminate other aspects of the host-pathogen interaction that have largely been unstudied in mechanistic studies. Here, we utilized a simple digital imaging platform to show that it is possible to identify the genetic basis of these other defense-related traits, such as the color and shape of the developing lesion in addition to the traditional size of the lesion. This shows that the generation of these visual traits is genetically determined by variation in the host, the pathogen and the interaction of genetic variation in the two organisms similarly to lesion size but can identify genes not known to influence lesion size.

Conducting GWAS within the *A. thaliana* accessions and mutant validation studies showed that these four traits had a blend of distinct and overlapping genetic mechanisms controlling their generation. Classical *B. cinerea* resistance genes, like *COI1* and *PAD3*, influenced both color and size of the developing lesion, but not the shape of the lesion, showing the role of known resistance mechanisms in these new lesion traits (Thomma et al., 1998; Ferrari et al., 2003; Kliebenstein et al., 2005; Rowe et al., 2010)(Figure 7). In contrast, some known resistance genes like *ACD6* affected only the color but not the size or shape of the lesion (Rate et al., 1999; Song et al., 2004; Lu et al., 2009)(Figure 7). More importantly, using new lesion-related traits for GWAS identified new genes involved in resistance to *B. cinerea* that had not previously been identified. This includes genes involved in light signal transduction, LUX, PHOT1 and PHYE, as well as sulfur transport (Clack et al., 1994; Devlin et al., 1998; Briggs and Christie, 2002; Halliday and Whitelam, 2003; Kataoka et al., 2004; Hazen et al., 2005; Zuber et al., 2010a; Nusinow et al., 2011; Sanchez-Lamas et al., 2016). The three light-related genes affected different lesion traits with PHOT1 being specific to lesion size, PHYE altering lesion size and yellowing while LUX and the sulfur transporter specifically altered lesion size (Figure 7). Thus, even these three genes that might have been expected to have overlapping effects show different and distinct effects from each other. This shows that there are specific mechanisms for each set of the traits and to fully understand the interaction requires more than simply measuring pathogen growth.

## Lesion Eccentricity as an Indicator of Pathogen Success

One difficulty of new traits from high-throughput digital imaging is to comprehend their possible biological role. For example lesion eccentricity (i.e. deviation from circularity) is a trait that is genetically controlled by both the pathogen and the host genes. Heritability of lesion eccentricity was 4.3% for plant accession, <0.1% for isolate, but jumped to 11.1% for accession:isolate, indicating that variation in this trait is highly dependent on the interaction of plant and pathogen genotypes. In two of the four pathogen genotypes, B05.10 and Supersteak, there was a negative correlation between lesion size and eccentricity suggesting that eccentric lesions tend to be smaller (Figure 3). In previous work, we had shown that eccentric lesions are associated with preferential growth along the primary vasculature, which implies that there is a shift from general radial outward growth to directed growth along the leaf midrib (Corwin et al., 2016a). Purely radial growth is typically considered to maximize local growth in microbial organisms and that would argue that directed growth may foresake potential local growth. A possible reason for forsaking local growth is that this could be a virulence strategy of some pathogen isolates to more rapidly move systemically through the vasculature of the plant. The fact that there are plant genes specific to affecting this trait indicates that *A. thaliana* has specific mechanisms that are geared towards altering the pattern of pathogen growth. As such, the success of the pathogen in the host-pathogen interaction would be a combination of the ability to infect the local tissue as well as move to distal tissue rapidly. Equally, if the pathogen has different growth strategies to optimize fitness, the plant would have to defend against local growth and growth geared towards spread of the pathogen.

## Lesion Color Genetic Mechanisms and Agriculture

It is harder to assess the potential ecological or evolutionary role of specific genetic mechanisms influencing the color of a lesion. However, lesion color is a potentially key aspect of the plant-pathogen interaction for vegetable and fruit crops. For example, a very small lesion that gives rise to a highly noticable color (e.g. Figure 1c) on a fruit or vegetable would greatly decrease the consumer acceptance of that product and lead to post-harvest crop loss if the lesion was large. As such, understanding the specific genetic mechanisms influencing the different color traits can have significant agricultural importance and they are presently not studied. Our study showed that as previously described, COI1 plays a key role in controlling the development of yellow in the plant-pathogen interaction (Rowe et al., 2010). This agrees with the known connection of jasmonate signaling to photosynthetic/plastid functioning (Kazan and Manners, 2011; Attaran et al., 2014; Campos et al., 2016). Interestingly, this also showed that additional genes not previously known to affect *B. cinerea* resistance also function in determining the level of lesion yellowness, like PHYE (Figure 8).

Genes associated with lesion greenness were enriched in activation of the immune system, activation of the innate immune system, and movement of cellular or subcellular components. These biological processes have obvious roles in plant response to a fungal infection, but their links with lesion greenness are less obvious. Previous studies had identified green islands in some plant-pathogen interactions, dead green tissue within pathogen lesions, as associated with increased levels of cytokinins and polyamines that delay senescence, as well as a variety of fungal toxins (Walters et al., 2008). We previously attempted to extract green color from lesions using both methanol and hexane, but were unsuccessful, indicating that the source of this greenness is not free chlorophyll (Corwin et al., 2016a). Further, greener lesions had no living plant cells from what we could assess with trypan blue staining (Corwin et al., 2016a). However, the genes associated with greenness included known *B cinerea* resistance genes like COI1, PAD3 and PAD4 (Figure 7). Interestingly, the only gene solely associated with lesion greenness was ACD6, a gene that is key to regulating plant cell death in response to pathogen attack. This suggests that there may be a role for how the plant cell dies in controlling this color. While our results do not provide an immediate direct mechanism for the color traits, it does show that there are potential mechanisms specific to the color traits in plant-pathogen interactions that maybe useful for understanding the infection process.

## Conclusion

This study has utilized high-throughput digital imaging of plant-pathogen interactions to uncover the genetics for previously uncharacterized lesion traits. Many of the candidate genes that were tested with insertional mutants affected more than one aspect of the plant-pathogen interaction, color, lesion size or shape of the lesion. But no gene affected all of these aspects of the interaction. As most studies of quantitative resistance focus on biomass or lesion size measurements while not studying lesion shape or color, our findings suggest that there are potentially unrecognized mechanisms that may be important for plant-pathogen interactions. Further validation studies could reveal more large-effect plant genes affecting quantitative resistance, filling gaps in our knowledge of cellular mechanisms involved in lesion traits and pathogen virulence and deserving of attention in plant breeding programs. Equally, these traits raise the need for broader life-history studies of plant-pathogen interactions in the field to identify the potential ecological or evolutionary drivers of these traits. GWA mapping coupled with high-throughput digital measurement of virulence-associated traits will be a useful tool in understanding broader plant-pathogen interactions.

## Materials & Methods

### Population Selection for GWAS

We used a previously published collection of 96 *A. thaliana* accessions that was chosen for pathogen resistance GWAS based on similarity in flowering time (63.1 ± 0.95 (s.e.) days to flowering). This population minimizes the indirect effects of ontogenic variation caused by a wide range of flowering times, while also reducing the effect of rare, medium effect alleles in order to inflate genetically-related residuals (Corwin et al., 2016b). Selection based on flowering time also removed genetic outliers and minimized population structure. The selected population extends broadly across the known phylogeny of *A. thaliana* accessions, increasing the resolution of the association study (Corwin et al., 2016b).

### Growth Conditions and Pathogen Infection

Seeds were cold stratified in 1% Phytagar at 4°C for seven days prior to sowing. Seeds were then sown in a randomized complete block design with three seeds per cell, into four 104-cell flats containing standard potting soil (Sunshine Mix #1, Sun Gro Horticulture, Agawam, MA). Plants were covered with a transparent humidity dome, placed in short day (10h full spectrum light) conditions in a controlled environment at 22°C, and watered biweekly as needed. After one week, seedlings were thinned to one seedling per pot. Sowing was repeated two weeks later to create another balanced randomized experiment so that eight biological replicates per accession were present across the two independent experiments in a randomized complete block design.

After five weeks of growth, mature leaves were excised, and placed on agar flats (Denby et al., 2004; Rowe and Kliebenstein, 2008). Individual leaves were inoculated with either one of four phenotypically and genotypically diverse isolates of *B cinerea* (Apple517, B05.10, Supersteak, UKRazz) or a mock-inoculated control (Atwell et al., 2015; Corwin et al., 2016b; Corwin et al., 2016a). Spores of five-day old sporulation cultures grown on organic peach slices were collected in sterilized 1/2x organic grape juice, counted with a hemocytometer, and standardized to a solution of 10 spores μl^−1^. Individual leaves were inoculated with a single 4μl droplet of one of the *B. cinerea* isolates or mock-inoculated with a control of 1/2x organic grape juice, for approximately 40 spores per leaf. The inoculated leaves were kept under transparent humidity domes at room temperature to allow lesions to grow for three days. At 72 hours post inoculation (hpi), photos were taken of infected leaves along with a 1 cm^2^ scale for size reference using an 18 Megapixel high resolution T3i Canon camera outfitted with an EF-S 10-22mm f/3.5–4.5 USM ultra-wide angle lens, achieving a resolution of approximately 10 pixels per mm (Corwin et al., 2016b).

### Image Analysis

A semi-automated image analysis script using the open-source R statistical environment (R Development Core Team, 2016) and the Bioconductor packages EBImage and CRImage (Failmezger H, 2010; Pau et al., 2010) was used to measure lesion traits such as lesion size, shape, and color (Corwin et al., 2016b). Briefly, the script identified leaves as objects that are have a green hue and highly saturated at their perimeter, whereas lesions were brown objects of low saturation within the leaf perimeter. Lesion and leaf mask images were generated and manually refined, or corrected where faulty. The area in pixels of these leaves and lesions, as well as the numbers and proportions of pixels of certain colors, and dimensions such as major and minor axes, perimeter, and eccentricity, were recorded. These values were converted to mm^2^ using a 1 cm reference standard contained within each image.

### Genome Wide Association Mapping

For each trait, we modeled heritability and obtained the model corrected least squared means using the linear model:

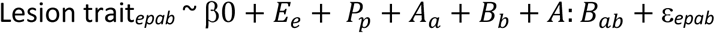

where E is experimental replicate, P is the individual plant, A is the genotype of the Arabidopsis accession and B is the genotype of the Botrytis isolate. Experimental replicate was modeled as a random effect, whereas individual plant, Arabidopsis accesion and Botrytis isolate were categorical. Publicly available data on genomic polymorphisms of the selected *A. thaliana* accessions was collected from the *A. thaliana* 1,001 genomes project (Ossowski et al., 2008; Cao et al., 2011; Gan et al., 2011; Schneeberger et al., 2011; Long et al., 2013; Alonso-Blanco et al., 2016). SNPs with minor allele frequency (MAF) < 0.2 were filtered out, resulting in a set of 115,310 SNPs for the 95 accessions. Association mapping was performed using the bigRR package in the R statistical environment (Shen et al., 2013). This previously published and validated method uses ridge regression, an appropriate approach given the small proportion of phenotypic variance attributable to genotype (Corwin et al., 2016b; Francisco et al., 2016). The ridge regression models the effects of all polymorphisms in a single model, treating each SNP as a random effect and introducing a bias to the regression estimates to reduce standard error. Thus, each polymorphism is assigned a heteroscedastic effect size (HEM), rather than a p-value, which is difficult to determine for random variables. Instead, a significant effect threshold value is delineated by permuting the phenotype data as it corresponds to the polymorphism data 1,000 times, and taking the 95^th^ and the 99^th^ percentiles. A gene is considered to be associated with a trait when two or more SNPs within the coding region have an effect size greater than the 95^th^ percentile threshold. This is a functional genome wide prediction method that has been shown to provide similar results to EMMA based GWAS (Kooke et al., 2016). Further, this GWAS pipeline has yielded a high rate of gene validation for a number of traits within *A. thaliana* (Corwin et al., 2016b; Francisco et al., 2016).

### GO Enrichment Analysis

Gene ontology (GO) enrichment analysis for biological processes was performed on all genes significantly associated with each of the four lesion traits using the Bioconductor packages org.At.tair.db, topGO, and goProfiles in the R statistical environment. Genes within *A. thaliana* that contain at least two significant SNPs that were associated with the trait of interest in the dataset were used as the genomic background sample for the analysis.

### Validation Tests

A selection of 23 *A. thaliana* genes that were found to be highly associated with Lesion Size, Lesion Greenness, Lesion Yellowness, and Lesion Eccentricity was made based on ready availability of viable homozygous T-DNA knockout lines. Seventeen of the lines had been previously identified and tested for lesion size and an additional six genes were chosen based on the magnitude of predicted effect size (Corwin et al., 2016b). Plants were grown as described above and no genotypes showed any evidence of flowering within this time frame in these conditions. At five weeks of age, the six first fully mature leaves were excised and inoculated with the four isolates of *B. cinerea* mentioned above at the same concentrations, and imaged at 72 hpi. These leaves showed no visual evidence of senescence at the start of the experiment and the control grape juice mock-inoculated leaves showed no developing lesion during the image analysis. The experiment was carried out in a randomized complete block design, with two experimental replicates each containing 10 biological replicates, amounting to 20 replicates for each *A. thaliana* mutant / *B. cinerea* isolate combination. Statistical differences between the WT Col-0 background and each mutant genotype were assessed using the linear model:

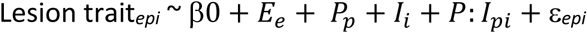

where E is experimental replicate, P is plant genotype, and I is the fungal isolate genotype. Experimental replicate was modeled as a random effect, where plant and fungal isolate genotypes were categorical.

## Acknowledgments

Financial support for this work was provided by the National Research Foundation DNRF grant 99, US NSF grants IOS 1339125, MCB 1330337 and IOS1021861, and the USDA National Institute of Food and Agriculture, Hatch project number CA-D-PLS-7033-H.

---

**Table S1. LSMeans for all measured lesion traits.** The model corrected mean value for all 75 measures lesion traits across all accessions for all isolates.

**Table S2. GO Enrichment Test using the 200 candidate genes associated with all four lesion Traits.**Annotated shows the total number of genes in the genome for the given term. Significant shows the number of candidate genes with this term and Expected is the number expected by random chance. Classic Fisher is the p-value for enrichment of this term.

**Table S3. Lesion Area GWA** Presented are the estimated effect sizes for all SNPs using lesion area for each isolate. The absolute value of the permutation estimated effect size are shown to the right for each isolate.

**Table S4. Lesion Greenness GWA** Presented are the estimated effect sizes for all SNPs using lesion greenness for each isolate. The absolute value of the permutation estimated effect size are shown to the right for each isolate.

**Table S5. Lesion Yellowness GWA** Presented are the estimated effect sizes for all SNPs using lesion yellowness for each isolate. The absolute value of the permutation estimated effect size are shown to the right for each isolate.

**Table S6. Lesion Eccentricity GWA** Presented are the estimated effect sizes for all SNPs using lesion eccentricity for each isolate. The absolute value of the permutation estimated effect size are shown to the right for each isolate.

**Figure S1:**
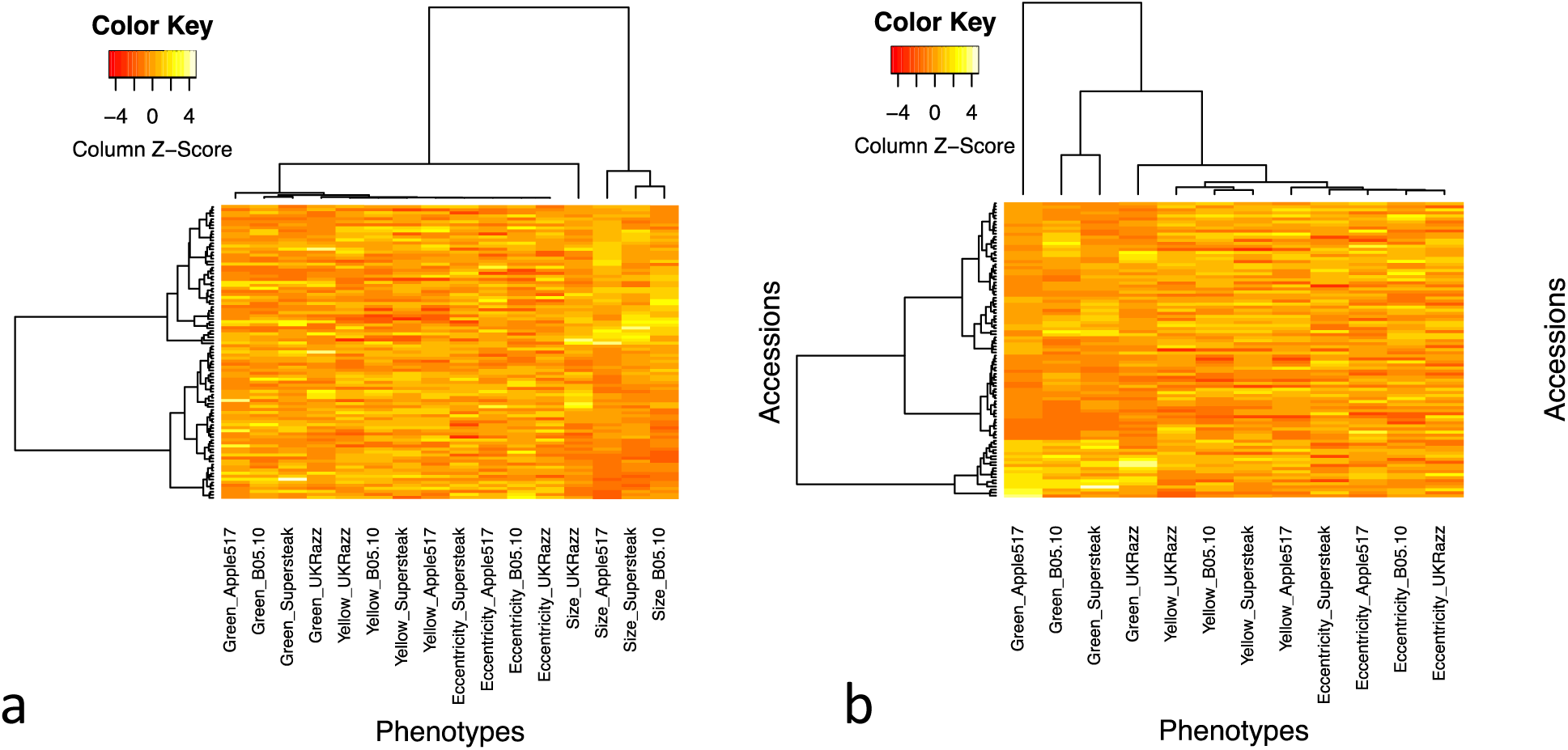
Hierarchical Cluster Analysis of Lesion Traits. Hierarchical cluster analysis of the a) four traits chosen to represent broad visual aspects of lesions, across the four pathogen isolates. b) Detail showing only color and shape traits illustrates relatedness and variation across accessions and isolates.

**Figure S2:**
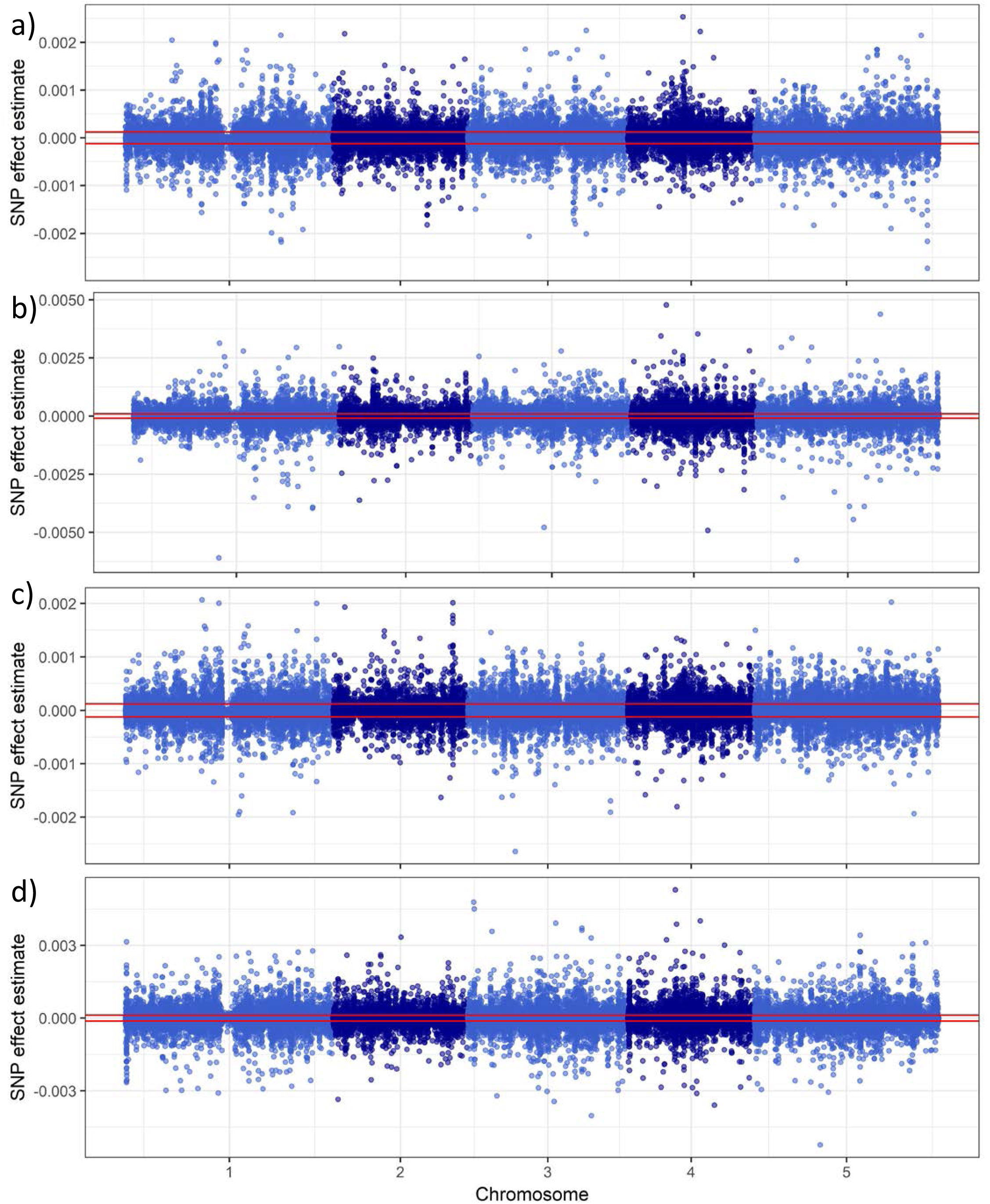
Lesion Greenness Manhattan Plots. Manhattan plots showing lesion greenness trait GWAS results, as measured on four *B. cinerea* isolates: a, Apple517; b, B05.10; c, Supersteak; and d, UKRazz. The horizontal red-line shows the significance threshold as estimated by permutation.

**Figure S3:**
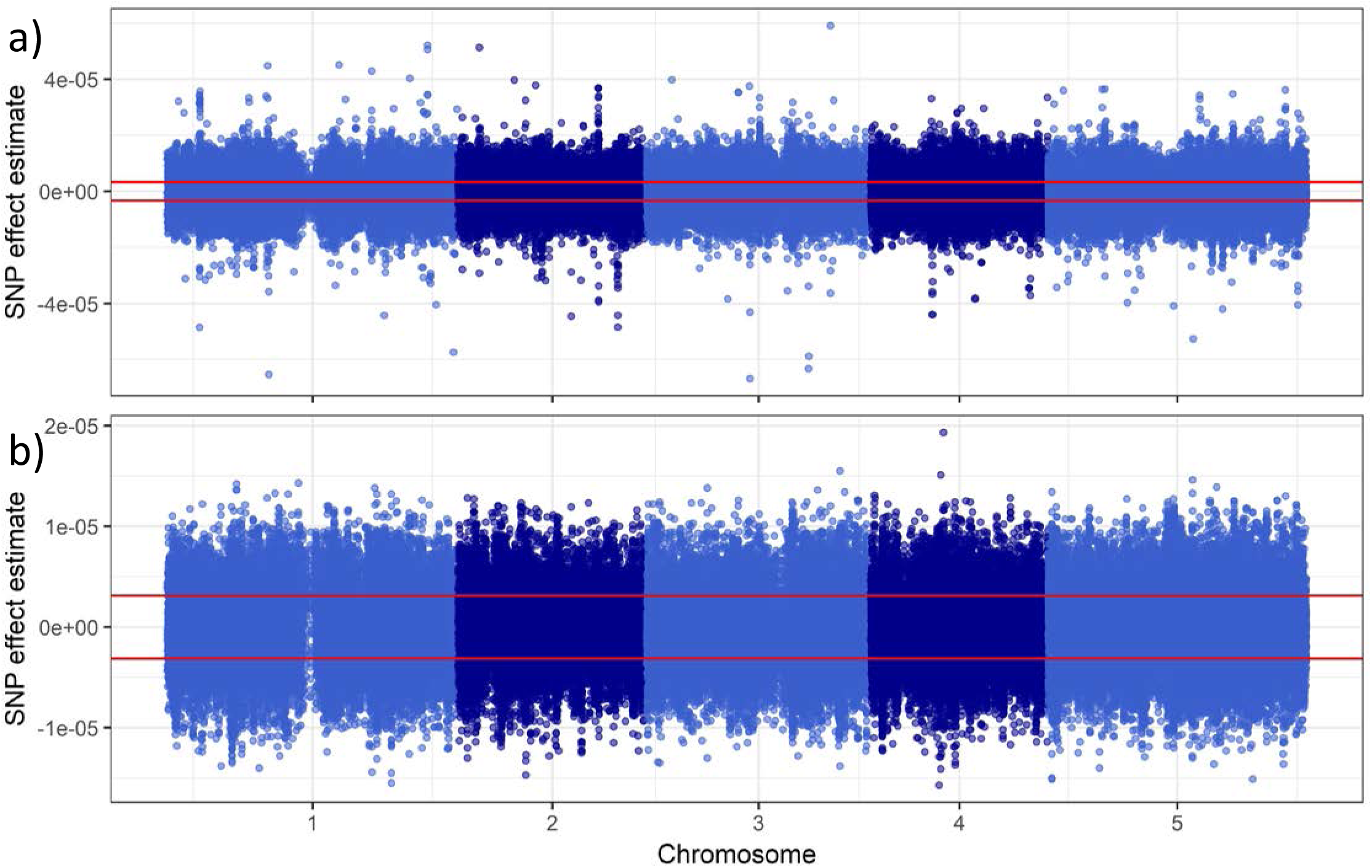
Lesion Yellowness and Eccentricity Shape Manhattan Plots for Apple517. Manhattan plots showing lesion yellowness (a) and eccentricity trait (b) GWAS results for Apple517. There were no significant SNPs identified for the other isolates. The horizontal red-line shows the significance threshold as estimated by permutation.

